# VF Mutations Lock JAK2 and TYK2 Kinase Dimers in a More Stable Conformation: Molecular Dynamics Reveals Enhanced Interface Rigidity

**DOI:** 10.1101/2025.05.16.654435

**Authors:** Tianwen Shao, Jiayi Yang, Yuxuan Cheng, Pingan Yuan, Haojie Cheng, Qi Yin, Jingyan Bai, Xiang Shen, Jianyu Zhu, Min Gan, Han Yang, Shengjie Sun

**Author notes:** Corresponding author: Han Yang Shengjie Sun. These authors contributed equally to this work (Tianwen Shao, Jiayi Yang, Yuxuan Cheng).

## Abstract

Homologous oncogenic valine-to-phenylalanine (VF) mutations, JAK2V617F and TYK2V678F, are key drivers of constitutive JAK/STAT signaling and pathogenesis of myeloproliferative neoplasms (MPNs) and rare TYK2-associated hematologic disorders. However, their atom-level mechanisms and divergent structural impacts remain poorly defined, hindering precision therapeutic development. Here, we conducted 4 × 100 ns all-atom molecular dynamics simulations to systematically compare dimer stability and interfacial interaction networks between wild-type and VF-mutant JAK2/TYK2. Both VF mutations significantly enhanced dimer stability by increasing the abundance and persistence of inter-monomer hydrogen bonds and salt bridges. The VF mutants also increased electrostatic complementarity at the PK–PK interface and induced novel stabilizing inter-chain contacts involving FERM domains. Consistent with this, MM/PBSA binding free energy calculations confirmed stronger energetic preference for dimerization in TYK2V678F, while JAK2V617F conferred only moderate stabilization. Collectively, our findings uncover how homologous oncogenic mutations drive distinct structural and energetic consequences across JAK paralogs, directly linking mutation-specific dimer stabilization to aberrant signaling in MPNs and rare hematologic diseases.

## Introduction

Janus kinases (JAKs) are a family of non-receptor tyrosine kinases, comprising JAK1, JAK2, JAK3, and TYK2.^1,2^ They play a central role in cytokine signaling through the JAK-STAT pathway.^3^ Upon cytokine binding to specific receptors, receptor dimerization occurs, leading to activation of associated JAKs. Activated JAKs phosphorylate receptor cytoplasmic tails, creating docking sites for STAT. Recruited STATs are subsequently phosphorylated, dimerize, and translocate to the nucleus, where they regulate the transcription.^4,5^ The JAK/STAT signaling pathway is indispensable for maintaining hematopoietic homeostasis, as it precisely governs the production, differentiation, and maturation of blood cells.^6^ Dysregulation of the JAK-STAT pathway is frequently associated with a wide range of diseases, including autoimmune disorders such as rheumatoid arthritis and hematologic malignancies.^7^ Naturally occurring mutations in cytokine receptors, JAKs, and STATs lead to immunodeficiency and myeloproliferative disorders in humans^8,9^. In particular, Gain-of-Function (GOF) mutations in JAK genes underlie a group of blood disorders known as myeloproliferative neoplasms (MPNs), including polycythemia vera, primary myelofibrosis, essential thrombocythemia, and leukemias.^10^

JAK family members share a conserved modular architecture consisting of seven JAK homology (JH) domains: an N-terminal four-point-one, ezrin, radixin, moesin (FERM) domain, an Src homology 2 (SH2) domain, and a C-terminal tandem kinase cassette composed of a regulatory pseudokinase (PK) domain and a catalytically active tyrosine kinase (TK) domain (Fig. 1).^11,12^ The TK domain mediates substrate phosphorylation, whereas the PK domain plays key autoinhibitory and regulatory roles^13,14^. A missense mutation in the PK domain of JAK2 (V617F) was shown to confer constitutive kinase activity and to be present in more than 90% of patients with polycythemia vera and approximately 50% of patients with essential thrombocythemia and primary myelofibrosis.^15-18^, Analogous mutations in other human JAK paralogs likewise result in constitutive activity, suggesting a shared activation mechanism across JAK family members, potentially involving ligand-independent dimerization at the cell surface.^19-21^ In 2022, a cryo-electron microscopy structure of full-length mouse JAK1 in complex with the intracellular Box1/Box2 regions of interferon λ receptor 1 (IFNλR1) was reported. This structure captured an activated JAK1 homodimer bearing the oncogenic V658F mutation, which corresponds to V617F in JAK2 (Fig. 1a) and V678F in TYK2 (Fig. 1b). The study proposed that the VF mutation enhances PK-domain dimerization by improving structural and hydrophobic complementarity at the PK–PK interface compared to the wild type.^22^

**Figure 1.**
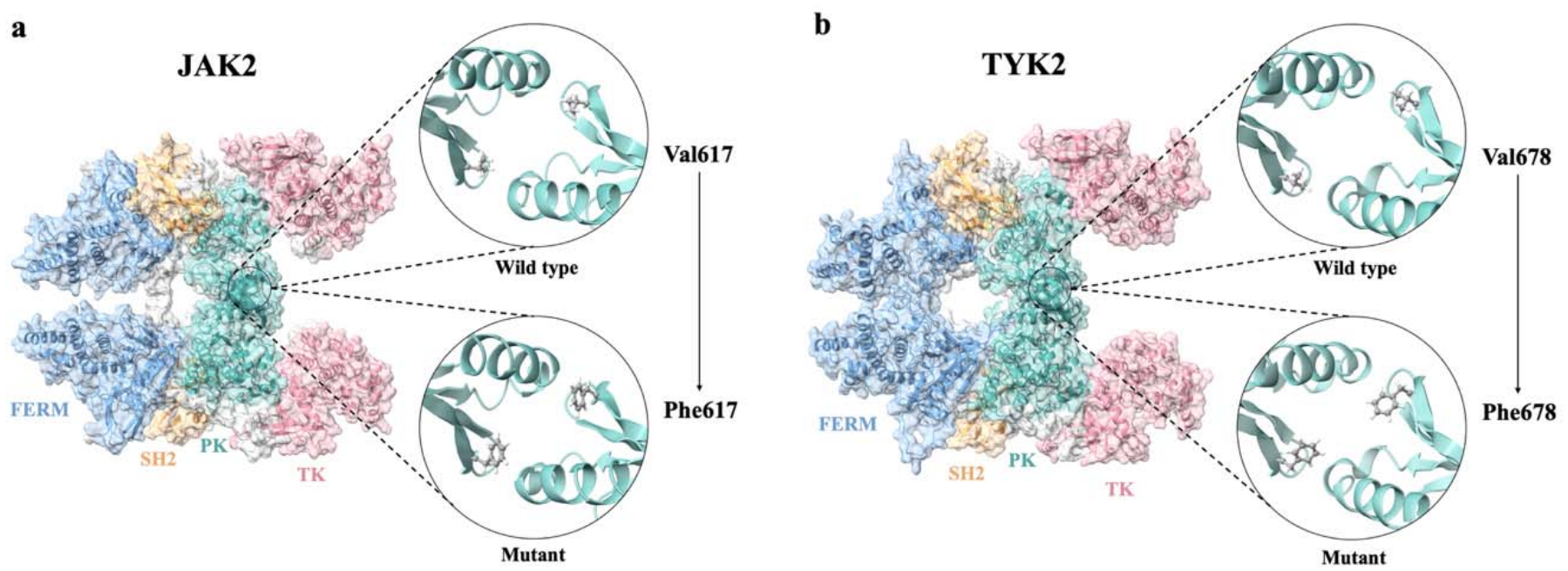
Domain organization and VF mutation sites in JAK2 and TYK2 dimers. (a) JAK2 and (b) TYK2. The FERM, SH2, pseudokinase (PK), and tyrosine kinase (TK) domains are colored blue, orange, teal, and pink, respectively; connecting regions are shown in gray. Enlarged views show the corresponding PK-domain regions in the wild-type and mutant structures, with Val617/Phe617 in JAK2 and Val678/Phe678 in TYK2 represented as ball-and-stick models in both monomers.

Despite these advances, experimental techniques remain limited in their ability to resolve detailed interactions at protein dimer interfaces, making it challenging to quantitatively characterize how VF mutations alter JAK dimerization. Advances in computational approaches have established molecular dynamics (MD) simulations as a powerful tool for probing mutation-induced structural and energetic changes at atomic resolution.^23^ Here, we employed all-atom MD simulations to investigate the atomic-level mechanisms by which the homologous JAK2V617F and TYK2V678F mutations differentially influence dimer stability and interfacial interactions. Our results provide a quantitative framework for understanding how VF mutations promote JAK dimerization and highlight distinct structural effects across JAK family members. These findings may further inform the design of selective therapeutics targeting mutant-specific JAK dimer interfaces while minimizing unintended activation of other JAK paralogs.

## Methods

### Modelling

Human JAK2 (UniProt ID: O60674) and TYK2 (UniProt ID: P29597) protein sequences were retrieved from the UniProt database.^24^ Homology models of full-length wild-type JAK2 and TYK2 were generated using SWISS-MODEL,^25^ employing the cryo-EM structure of full-length mouse JAK1 (PDB ID: 7T6F) as the template. The oncogenic mutations (V617F in JAK2 and V678F in TYK2, analogous positions in the PK domain) were introduced using the Dunbrack rotamer library^26^ in Chimera to select the most probable side-chain conformation.

Each dimer was placed in a rectangular simulation box and solvated with TIP3P water molecules,^27^ maintaining a minimum distance of 10 Å from the protein to the box edge. The systems were ionized with 150 mM NaCl to achieve electrostatic neutrality. All simulations employed the CHARMM36m force field.^28^ The composition and initial box dimensions of each simulation system are provided in Table S1 of the Supporting Information. A representative solvated simulation box is shown in Fig. S1.

### Molecular dynamics simulation

All MD simulations were carried out using NAMD3.0.^29^ Each system was first energy-minimized for 20,000 steps using the conjugate gradient algorithm, followed by a 100 ns unrestrained NPT production run. A time step of 2 fs was used. The temperature was set to 310.15 K, and the pressure was set to 1 atm using a Langevin thermostat with a damping coefficient of 1 ps□^1^ and Nosé–Hoover Langevin piston barostat with a decay period of 50 fs, respectively. Periodic boundary conditions were applied to the solvated simulation box. Long-range electrostatic interactions were calculated using the particle-mesh Ewald (PME) method. Short-range nonbonded interactions were truncated with a cutoff distance of 12 Å, and a switching function was applied starting at 10 Å. Trajectory frames were saved every 100 ps for analysis.

### Root-mean-square deviation (RMSD) and Root-mean-square fluctuation (RMSF)

RMSD of the backbone atoms was calculated using VMD1.9.3,^30^ to assess simulation stability (Equation 1),

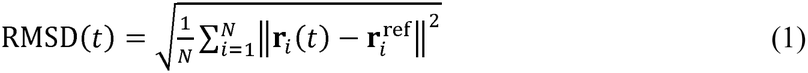

where *N* is the total number of backbone atoms, *r*_*i*_ represents the position of atom *i* at time *t* after least-squares alignment to the reference structure, and 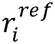 denotes the corresponding position of atom *i* in the reference structure. The initial structure was used as the reference. RMSF of Cα atoms was calculated for each residue to evaluate local flexibility (Equation 2),

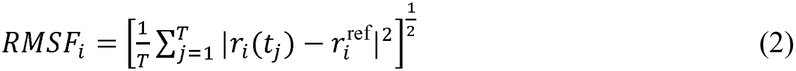

where *T* represents the number of frames taken from MD simulation time, and *i* represents the residue ID. *r*_*i*_ (*t*_*j*_) is the position of residue *i* at time *t*_*j*_. The 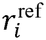 is the reference position (time-averaged position) of residue *i*.

### Hydrogen bonds and salt bridges

Inter-monomer hydrogen bonds and salt bridges were analyzed using the Hydrogen Bonds and Salt Bridges plugins in VMD 1.9.3, respectively. Hydrogen bonds were identified using a donor– acceptor heavy-atom distance cutoff of 4.0 Å and an angle cutoff of 20°. The two monomers were specified as separate atom selections, with both monomers allowed to act as hydrogen bond donors and acceptors. Inter-monomer salt bridges were identified using a distance cutoff of 4.0 Å between the side-chain oxygen atoms of acidic residues and the side-chain nitrogen atoms of basic residues.

### Electrostatic potential analysis

The electrostatic potentials were calculated using Delphi8.4.5.^31^ The charge and radius were assigned by pdb2pqr^32^ from CHARMM36m.^28^ The dielectric constants for the protein and water were set as 2 and 80, respectively. Salt concentration, probe radius, filling ratio of the protein, and resolution were set as 150 mM, 1.4 Å, 0.70, and 2 grid/Å, respectively. The resulting potential was mapped on the molecular surface using Chimera1.19,^33^ with surface colored red, blue and white, representing negative, positive and neutral, respectively.

### Electrostatic force

The electrostatic force between the PK domains of the two monomers was calculated using DelphiForce.^34^ After 100-ns MD simulations for each of the four models, the last frames of PK domains were extracted from trajectories. Within each model, the separation between the PK domains was increased from 5 to 20 Å in 1 Å increments. The binding force was calculated using Equation 3.

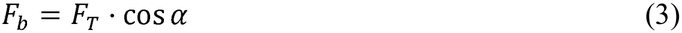

*F*_*T*_ and *F*_*b*_ represent the total and binding force, respectively, while α represents the angular deviation between the electrostatic force and the line connecting the two mass centers.

### MM/PBSA binding free energy

For each of the four systems, the last 20 ns of the production trajectory were used for analysis. Two monomers were separated from dimer to calculate the binding free energy for the dimer.

The free energy Δ*E*_*f*_ was calculated by the sum of the Coulombic, VDW, polar, and nonpolar solvation binding energies (Equation 4),

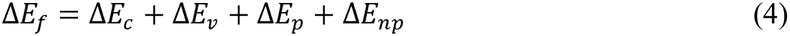

where Δ*Ec*, Δ*E*_*v*_, Δ*E*_*p*_, and Δ*E*_*np*_ represent the Coulombic binding energy, van der Waals (vdW) binding energy, polar solvation energy, and nonpolar solvation energy, respectively. The Coulombic energy describes direct electrostatic interactions between solute atomic charges, the vdW energy accounts for repulsive and dispersion interactions between solute atoms, the polar solvation energy represents the contribution from solvent dielectric polarization, and the nonpolar solvation energy approximates nonpolar solvent effects using a linear function of the solvent-accessible surface area (SASA). The Coulombic binding energy and polar solvation energy were calculated using DelPhi, while the vdW binding energy was calculated using NAMD3. The nonpolar solvation energy was calculated from SASA according to Equation 5:

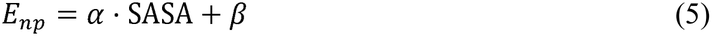

Here, α is the surface tension coefficient, and β is the constant offset used to estimate the nonpolar solvation energy from SASA.

The binding energy (Δ*E*) for each component was calculated using Equation 6:

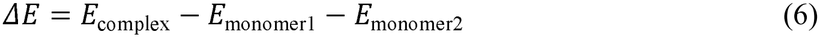

## Results

### System stability and simulation convergence

Backbone RMSD profiles over the 100 ns trajectories are shown in Fig. 2a. The VF-mutant systems (JAK2V617F and TYK2V678F) exhibited earlier RMSD plateaus than their corresponding wild-type dimers.

**Figure 2.**
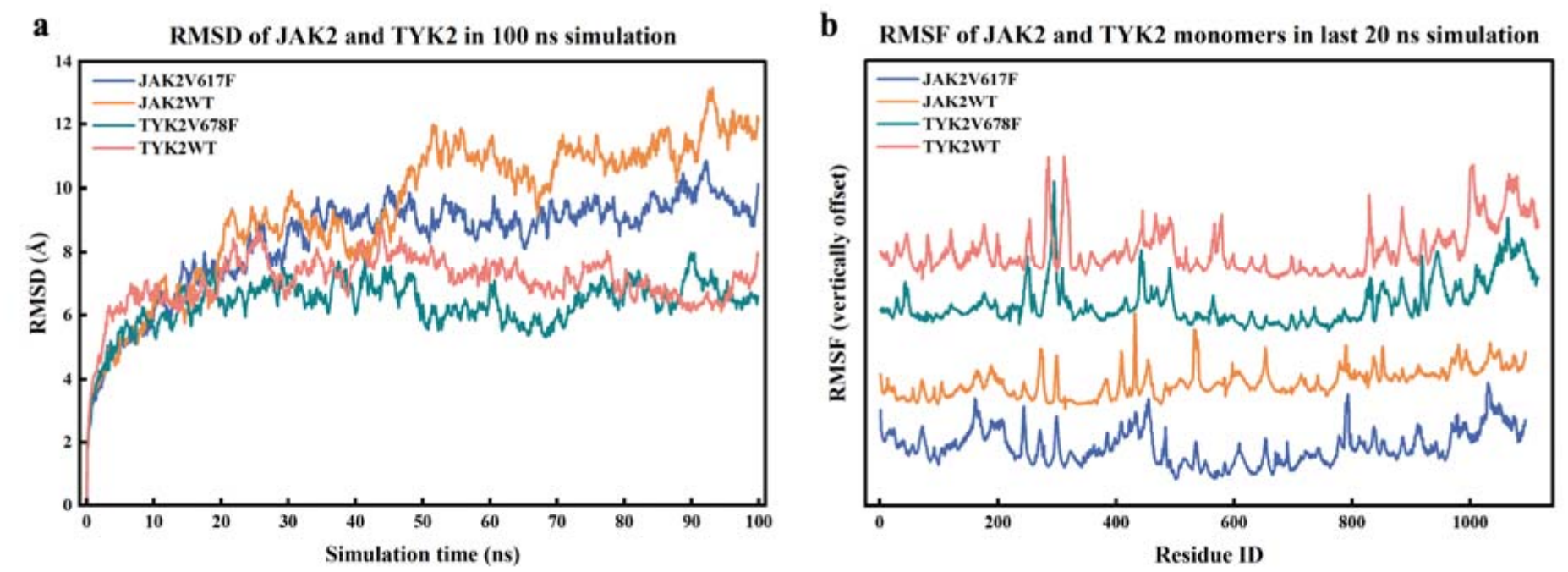
RMSD and RMSF analysis of JAK dimers. (a) Backbone RMSD of the four systems as a function of simulation time over 100 ns. (b) Cα RMSF of JAK2 and TYK2 monomers over the last 20 ns of each trajectory. The RMSF profiles were vertically offset for clarity; their relative vertical positions do not represent differences in absolute RMSF values.

RMSF analysis further revealed reduced flexibility at the PK domain dimer interface in both VF-mutant systems relative to wild type (Fig. 2b). This decrease in local flexibility indicates increased rigidity of the dimerization interface, consistent with a stabilizing effect of the VF mutations.

### VF mutation enhanced the hydrogen bonds and salt bridges in both JAK2 and TYK2 dimers

Inter-monomer hydrogen bonds and salt bridges were monitored over the last 20 ns of each trajectory (Figs. 3 and 4). Occupancy (%) represents the fraction of frames in which the interaction was present. Only hydrogen bonds with occupancies greater than 25% and salt bridges with occupancies greater than 10% are shown in the bar graphs. Complete occupancy data are provided in Tables S2–S5 of the Supporting Information.

**Figure 3.**
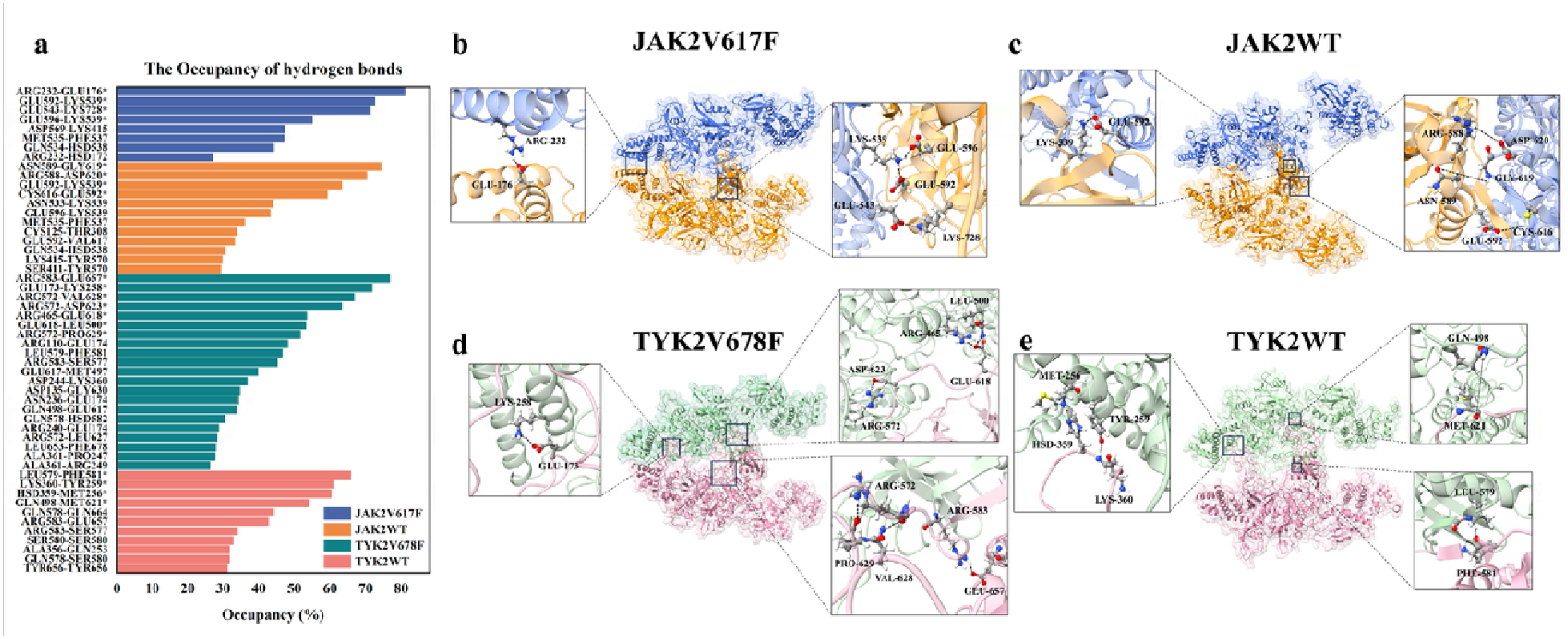
Inter-monomer hydrogen bond analysis of JAK dimers. (a) Bar plots of hydrogen bond occupancies for all four systems. Only hydrogen bonds with occupancies greater than 25% are shown; those exceeding 50% are considered stable and are marked with asterisks and highlighted on the corresponding structures shown on the right. (b) Stable hydrogen bonds in JAK2V617F. (c) Stable hydrogen bonds in JAK2WT. (d) Stable hydrogen bonds in TYK2V678F. (e) Stable hydrogen bonds in TYK2WT.

**Figure 4.**
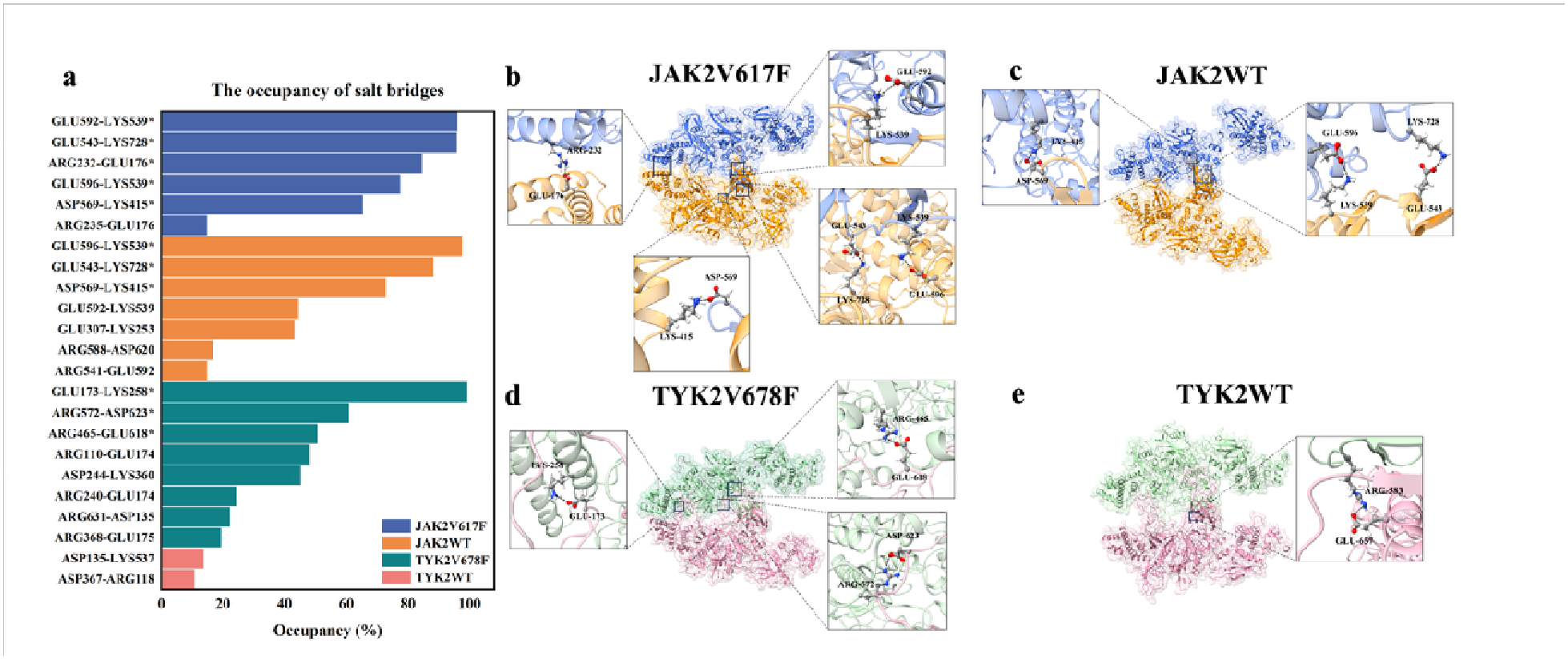
Inter-monomer salt bridge analysis of JAK dimers. (a) Bar plots of salt bridge occupancies for all four systems. Only salt bridges with occupancies greater than 10% are shown; those exceeding 50% are considered stable and are marked with asterisks and highlighted on the corresponding structures shown on the right. (b) Stable salt bridges in JAK2V617F. (c) Stable salt bridges in JAK2WT. (d) Stable salt bridges in TYK2V678F. (e) The low-occupancy Arg583–Glu657 salt bridge in TYK2WT, additionally shown for comparison despite its occupancy below 10%.

#### Hydrogen Bonds

In JAK2WT, stable hydrogen bonds were primarily confined to the PK domain. In contrast, JAK2V617F not only preserved these PK-domain interactions but also formed persistent new hydrogen bonds involving the FERM domain. The most persistent hydrogen bond in JAK2V617F was Arg232–Glu176, located in the FERM domain, with an occupancy of 81.25%; this interaction was absent in JAK2WT. Two prominent PK-domain hydrogen bonds, Glu592– Lys539 and Glu596–Lys539, were also more persistent in the mutant (occupancies of 72.75% and 55.25%, respectively) than in the wild-type system (63.5% and 43.5%). Conversely, several relatively stable hydrogen bonds observed in JAK2WT, such as Asn589–Gly619 (74.5%) and Arg588–Asp620 (70.5%), were not detected in JAK2V617F. While these wild-type interactions exhibited moderate stability, their occupancies were lower than those of the most persistent hydrogen bonds formed in the mutant (Fig. 3a-c).

Both TYK2WT and TYK2V678F formed stable hydrogen bonds within the PK and FERM domains. However, the specific interaction networks differed markedly. The V678F mutation led to a near-complete redistribution of high-occupancy hydrogen bonds. For example, in the FERM domain, TYK2V678F formed a prominent Glu173–Lys258 hydrogen bond (occupancy 72%), whereas TYK2WT exhibited interactions such as Lys360–Tyr259 (61%) and His359–Met256 (60.5%). In the PK domain, TYK2V678F showed strong hydrogen bonds including Arg583– Glu657 (77%) and Arg572–Val628 (67%), while TYK2WT predominantly formed Leu579– Phe581 (66%) and Gln498–Met621 (54.5%) (Fig. 3a-e).

#### Salt bridges

The VF mutations markedly enhanced the stability of inter-monomer salt bridges, with the effect being particularly pronounced in TYK2. TYK2V678F formed multiple persistent salt bridges, whereas TYK2WT exhibited virtually no high-occupancy salt bridge interactions. In TYK2WT, only a weak salt bridge between Arg583 and Glu657 was observed (Fig. 4a-e). Although the inter-residue distance fell below 4 Å in only 8% of frames, it fluctuated around 5.6 Å throughout the last 20 ns and was therefore classified as a weak interaction (Fig. S5), which is highlighted on the structure (Fig. 4e). Distance traces for the salt-bridge residue pairs in JAK2V617F, JAK2WT, TYK2V678F, and TYK2WT are provided in Figs. S2–S5, respectively.

In JAK2, the effects of the mutation were more heterogeneous. The salt bridges Glu592–Lys539 and Glu543–Lys728 were substantially more stable in JAK2V617F, with occupancies of 96.25% and 96%, respectively, compared with 44.75% and 88.5% in JAK2WT. In contrast, Glu596– Lys539 and Asp569–Lys415 were slightly more persistent in the wild-type system (97.75% and 73%) than in the mutant (77.75% and 65.5%) (Fig. 4a-c).

Notably, stable salt bridges involving the FERM domain emerged in both JAK2V617F and TYK2V678F, whereas no such interactions were observed in the corresponding wild-type systems.

### VF mutation enhanced the electrostatic interactions in both JAK2 and TYK2 dimers

Electrostatic surface potentials at the PK domain dimer interfaces are shown in Fig. 5a-d. In the JAK2WT system, the opposing interfaces were predominantly enriched with positive charges, whereas in the TYK2WT system the interfaces were mainly negatively charged. In contrast, both VF mutants exhibited clear electrostatic complementarity, with oppositely charged regions facing each other across the PK–PK interface.

Electrostatic binding force calculations revealed electrostatic attraction between the PK domains. The magnitude of the attractive force decreased with increasing separation distance and approached zero at larger distances, consistent with physical expectations. Compared with their respective wild-type dimers, JAK2V617F showed a modest increase in the calculated electrostatic attraction, whereas the increase was more pronounced in TYK2V678F (Fig. 5e).

**Figure 5.**
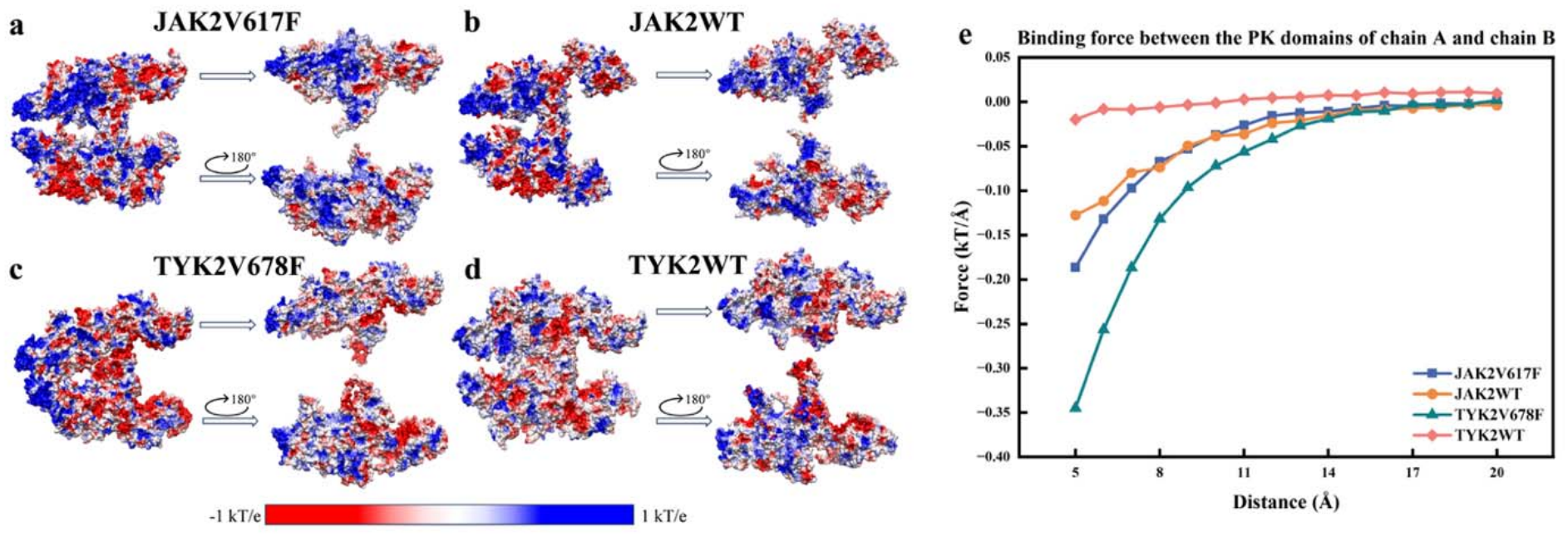
Electrostatic surface potential and inter-domain electrostatic forces of JAK dimers. (a) Electrostatic surface potential of JAK2V617F. (b) Electrostatic surface potential of JAK2WT. (c) Electrostatic surface potential of TYK2V678F. (d) Electrostatic surface potential of TYK2WT. (e) Electrostatic binding force between the PK domains as a function of inter-domain separation (5–20 Å, 1 Å increments) for all four systems.

### VF mutation decreased the binding free energy in both JAK2 and TYK2 dimers

MM/PBSA binding free energy calculations further quantified the effects of the VF mutations on dimer stability. In JAK2, the V617F mutation resulted in a modest improvement in binding free energy, with ΔE decreasing from –94.55 kcal/mol in the wild-type to -100.14 kcal/mol in the mutant. In contrast, the effect was markedly larger in TYK2, where the V678F mutation yielded a ΔE of –194.76 kcal/mol compared to –83.39 kcal/mol in the wild-type, corresponding to an improvement of more than 110 kcal/mol. These results indicate a substantially stronger energetic preference for dimer formation in TYK2V678F relative to JAK2V617F (Fig. 6a).

**Figure 6.**
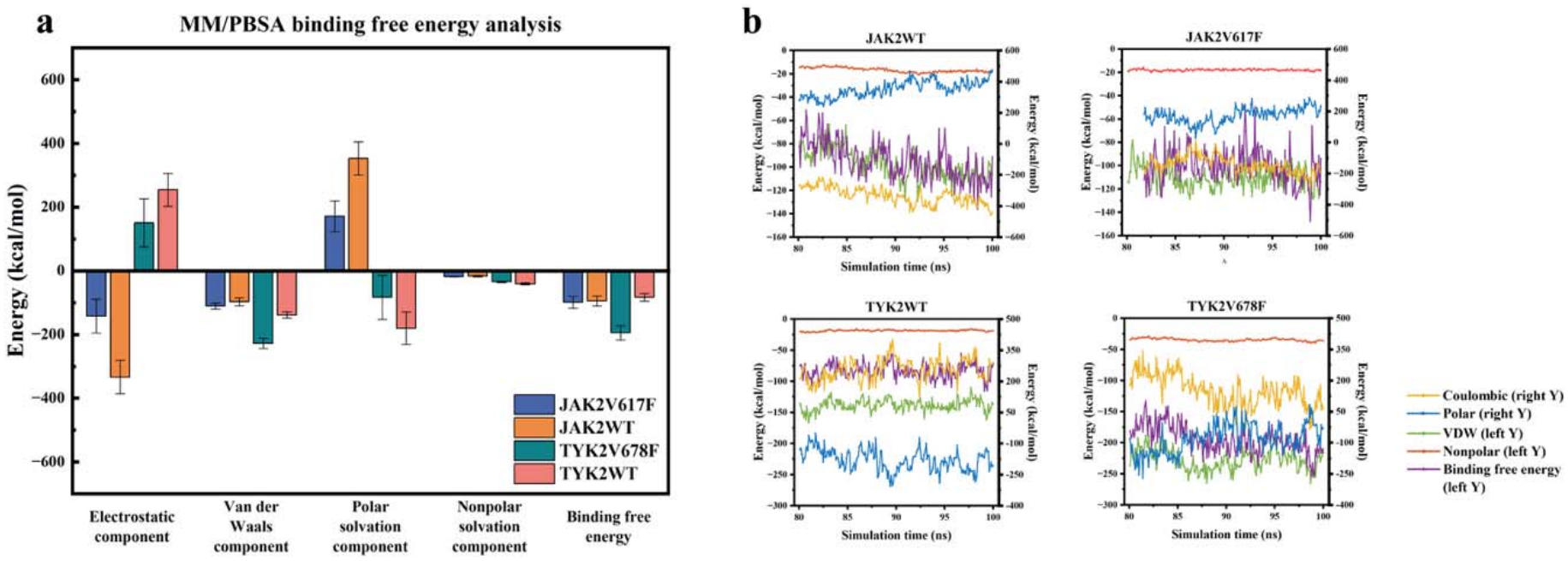
MM/PBSA binding free energy analysis of JAK dimers. (a) Bar plots show the contributions of individual binding free energy components for each system. From left to right, the components correspond to the electrostatic, Van der Waals, polar solvation, and nonpolar solvation terms. The rightmost bar represents the total binding free energy, calculated as the sum of all components. (b) Time evolution of each energy component for each system over the last 20 ns of the simulation.

Analysis of individual energy components indicates that the difference in binding free energy between JAK2V617F and JAK2WT mainly arises from a reduction in the polar solvation term, with a minor contribution from a decrease in the Van der Waals component. In contrast, for TYK2V678F, the difference relative to TYK2WT is dominated by the Van der Waals term, with an additional contribution from a modest decrease in the Coulombic component (Fig. 6b).

## Discussion

In this study, we employed all-atom molecular dynamics simulations to investigate how homologous VF mutations influence the stability of JAK2 and TYK2 dimers. Multiple complementary analyses consistently demonstrate that both JAK2V617F and TYK2V678F enhance dimer stability relative to their wild-type counterparts, with the stabilizing effect being substantially stronger in TYK2V678F.

These mutations reinforce hydrogen bonds, salt bridges, and electrostatic complementarity at the PK-domain interface. Notably, these stabilizing effects are not confined to the PK domain. In both mutants, persistent inter-monomer interactions involving the FERM domains emerge, whereas such interactions are absent or only transient in the corresponding wild-type dimers. This observation suggests that enhanced PK-domain dimerization may indirectly constrain the relative orientations and dynamics of distal domains, thereby promoting secondary inter-domain contacts. Such long-range, mutation-induced allosteric effects may contribute to the overall stabilization of the full-length JAK dimer.

MM/PBSA binding free energy analysis shows that both VF mutants exhibit lower binding free energies than their wild-type counterparts, with TYK2V678F displaying a markedly lower binding free energy than JAK2V617F. Decomposition of the binding free energy indicates that, relative to JAK2WT, the contribution to the binding free energy difference in JAK2V617F mainly comes from a reduction in the polar solvation term, with a slight contribution from the Van der Waals component. In contrast, for TYK2V678F, the binding free energy difference relative to TYK2WT is dominated by the Van der Waals component, accompanied by a modest decrease in the Coulombic term.

Despite the enhanced dimer stability observed in our simulations, it is important to emphasize that increased dimerization alone is unlikely to fully account for constitutive activation of the JAK-STAT signaling pathway. VF mutations may also promote signaling by relieving autoinhibitory interactions within individual JAK monomers.^22^ Moreover, enhanced structural stability does not necessarily translate directly into increased downstream signaling output. Consistent with this notion, previous studies have reported that TYK2V678F exhibits a much weaker effect on downstream STAT activation and cell proliferation than JAK2V617F,^35^ despite their structural homology. This discrepancy highlights the complexity of JAK regulation and suggests that additional factors beyond dimer stability—such as domain-specific regulatory interactions and cellular context—play critical roles in determining signaling outcomes.

Taken together, our results provide molecular-level insight into how VF mutations enhance JAK dimer stability through reinforced interfacial interactions, electrostatic complementarity, and long-range domain coupling. At the same time, the observed differences between JAK2 and TYK2 underscore that homologous oncogenic mutations can give rise to distinct structural and functional consequences across JAK family members. These findings help refine current models of JAK activation and may inform future efforts to design selective therapeutics targeting mutant-specific dimer interfaces while minimizing unintended activation of other JAK paralogs.

## Data Availability

This study was carried out using publicly available data from Protein Data Bank at https://www.rcsb.org/. The data supporting this article have been included as part of the Supplementary Information.

## Authors’ contributions

T.S., J.Y. and H.Y. drafted and wrote the main manuscript text. T.S., J.Y., P.Y., and H. C. performed the molecular dynamics (MD) simulations. J.Y., T.S., J.B. and Q.Y. carried out the data analyses and interpretation. X.S., J. Z., M.G., H.Y., and S.S. acquired the funding and provided supervision. All authors read and approved the final manuscript.

## Consent to publish

Consent to Publish declaration: not applicable.

## Ethics approval

Ethics declaration: not applicable.

