## Supplementary material for "VF Mutations Lock JAK2 and TYK2 Kinase Dimers in a More Stable Conformation: Molecular Dynamics Reveals Enhanced Interface Rigidity": SI

Corresponding author:

| **Table S1. Composition and initial box dimensions of the MD simulation systems.** | | | | |
| --- | --- | --- | --- | --- |
| **System** | **JAK2WT** | **JAK2V617F** | **TYK2WT** | **TYK2V678F** |
| Protein residues | 2,186 | 2,186 | 2,230 | 2,230 |
| Protein atoms | 35,488 | 35,496 | 35,230 | 35,238 |
| Water molecules | 87,290 | 87,173 | 81,540 | 81,536 |
| Na⁺ ions | 250 | 250 | 250 | 250 |
| Cl⁻ ions | 246 | 246 | 230 | 230 |
| Total atoms | 297,854 | 297,511 | 280,330 | 280,326 |
| Initial box length a (Å) | 115.655 | 115.192 | 116.210 | 116.210 |
| Initial box length b (Å) | 158.014 | 158.524 | 149.171 | 149.171 |
| Initial box length c (Å) | 175.849 | 175.470 | 175.032 | 175.032 |

Protein residue and atom counts refer to the complete dimer. Box dimensions refer to the initial solvated systems; all box angles are 90°.

| **Table S2. The details of interchain hydrogen bonds of JAK2WT and JAK2V617F.** | | | | | |
| --- | --- | --- | --- | --- | --- |
| **JAK2 WT Dimer** | | | **JAK2 V617F Dimer** | | |
| **Donor** | **Acceptor** | **Occupancy** | **Donor** | **Acceptor** | **Occupancy** |
| ASN589 | GLY619 | 74.50% | ARG232 | GLU176 | 81.25% |
| ARG588 | ASP620 | 70.50% | GLU592 | LYS539 | 72.75% |
| GLU592 | LYS539 | 63.50% | GLU543 | LYS728 | 71.50% |
| CYS616 | GLU592 | 59.50% | GLU596 | LYS539 | 55.25% |
| ASN533 | LYS539 | 44.33% | ASP569 | LYS415 | 47.50% |
| GLU596 | LYS539 | 43.50% | MET535 | PHE537 | 47.38% |
| MET535 | PHE537 | 36.30% | GLN534 | HSD538 | 44.50% |
| CYS125 | THR308 | 34.25% | ARG232 | HSD172 | 27.50% |
| GLU592 | VAL617 | 33.50% | ARG588 | ASP620 | 24.50% |
| GLN534 | HSD538 | 31.00% | ARG235 | GLU176 | 21.00% |
| LYS415 | TYR570 | 30.00% | ARG228 | ARG228 | 19.00% |
| SER411 | TYR570 | 29.50% | HSD172 | TYR231 | 18.00% |
| ARG588 | VAL617 | 21.00% | LYS415 | TYR570 | 17.00% |
| GLU543 | LYS728 | 21.00% | ASN542 | GLU592 | 16.00% |
| LYS539 | MET532 | 18.00% | ASN589 | GLY619 | 16.00% |
| ARG541 | ASN729 | 16.00% | ARG541 | GLU592 | 15.50% |
| LYS539 | MET535 | 16.00% | GLN239 | GLU173 | 13.50% |
| ASN589 | ASP620 | 15.00% | GLN534 | PHE537 | 12.50% |
| MET535 | VAL536 | 13.50% | MET535 | VAL536 | 12.00% |
| ASP313 | GLY516 | 13.00% | GLU592 | PHE617 | 10.50% |
| ASN533 | HSD538 | 12.00% |  |  |  |
| LYS414 | TYR570 | 11.50% |  |  |  |
| VAL536 | VAL536 | 11.50% |  |  |  |
| ASN589 | VAL617 | 10.50% |  |  |  |

| **Table S3. The details of interchain hydrogen bonds of TYK2WT and TYK2V678F.** | | | | | |
| --- | --- | --- | --- | --- | --- |
| **TYK2 WT Dimer** | | | **TYK2 V678F Dimer** | | |
| **Donor** | **Acceptor** | **Occupancy** | **Donor** | **Acceptor** | **Occupancy** |
| LEU579 | PHE581 | 66.00% | ARG583 | GLU657 | 77.00% |
| LYS360 | TYR259 | 61.00% | GLU173 | LYS258 | 72.00% |
| HSD359 | MET256 | 60.50% | ARG572 | VAL628 | 67.00% |
| GLN498 | MET621 | 54.50% | ARG572 | ASP623 | 63.50% |
| GLN578 | GLN664 | 44.50 % | ARG465 | GLU618 | 53.75% |
| ARG583 | GLU657 | 43.00% | GLU618 | LEU500 | 53.50% |
| ARG583 | SER577 | 34.33% | ARG572 | PRO629 | 52.00% |
| SER580 | SER580 | 33.00% | ARG110 | GLU174 | 48.50% |
| ALA356 | GLN253 | 32.00% | LEU579 | PHE581 | 46.90% |
| GLN578 | SER580 | 32.00% | ARG583 | SER577 | 45.50% |
| TYR656 | TYR656 | 27.50% | GLU617 | MET497 | 40.00% |
| ASP650 | GLY680 | 27.50% | ASP244 | LYS360 | 37.00% |
| ASP367 | ARG118 | 22.00% | ASP135 | GLY630 | 35.00% |
| HSD649 | SER647 | 22.00% | ASN236 | GLU174 | 34.50% |
| LEU653 | VAL678 | 21.00% | GLN498 | GLU617 | 34.00% |
| LEU579 | SER580 | 20.75% | GLN578 | HSD582 | 30.75% |
| GLN578 | LEU653 | 19.00% | ARG240 | GLU174 | 29.00% |
| HSD649 | PRO681 | 15.21% | ARG572 | LEU627 | 28.50% |
| GLN492 | MET621 | 14.50% | LEU653 | PHE678 | 28.25% |
| LYS360 | MET115 | 13.50% | ALA361 | PRO247 | 28.00% |
| ARG465 | GLU618 | 12.50% | ALA361 | ARG249 | 26.50% |
| ASP650 | VAL678 | 12.00% | ARG368 | GLU175 | 25.00% |
| GLN252 | LYS357 | 12.00% | GLN252 | GLN364 | 25.00% |
| HSD359 | TYR259 | 10.5% | GLU618 | SER499 | 25.00% |
|  |  |  | ARG583 | LEU579 | 23.50% |
|  |  |  | ARG631 | ASP135 | 23.50% |
|  |  |  | HSD649 | PRO681 | 23.25% |
|  |  |  | ARG233 | GLU174 | 21.50% |
|  |  |  | ARG490 | GLU618 | 21.00% |
|  |  |  | ARG118 | GLU371 | 20.50% |
|  |  |  | ARG233 | HSD177 | 20.50% |
|  |  |  | GLN364 | SER251 | 20.50% |
|  |  |  | SER580 | SER580 | 20.50% |
|  |  |  | ARG269 | GLU174 | 18.50% |
|  |  |  | GLN498 | GLU618 | 18.00% |
|  |  |  | ARG233 | GLU173 | 16.50% |
|  |  |  | ARG265 | GLU173 | 16.00% |
|  |  |  | LEU574 | PRO626 | 16.00% |
|  |  |  | ARG679 | ASP650 | 15.50% |
|  |  |  | ASP135 | PRO629 | 15.50% |
|  |  |  | LEU579 | SER580 | 15.00% |
|  |  |  | ALA366 | ASN116 | 14.50% |
|  |  |  | GLN253 | GLN364 | 14.00% |
|  |  |  | GLY363 | SER251 | 14.00% |
|  |  |  | GLN246 | HSD359 | 13.50% |
|  |  |  | GLY363 | LEU250 | 13.50% |
|  |  |  | ARG118 | GLU173 | 12.50% |
|  |  |  | ARG583 | LEU653 | 12.25% |
|  |  |  | ARG118 | GLU174 | 12.00% |
|  |  |  | GLY680 | HSD649 | 12.00% |
|  |  |  | ARG118 | GLU175 | 11.50% |
|  |  |  | HSD359 | PRO247 | 11.00% |
|  |  |  | ARG572 | GLU624 | 10.00% |
|  |  |  | ARG631 | SER133 | 10.00% |
|  |  |  | HSD582 | SER577 | 10.00% |

| **Table S4. The details of interchain salt bridges of JAK2WT and JAK2V617F.** | | | | | |
| --- | --- | --- | --- | --- | --- |
| **JAK2 WT Dimer** | | | **JAK2 V617F Dimer** | | |
| **Residue1** | **Residue2** | **Occupancy** | **Residue1** | **Residue2** | **Occupancy** |
| GLU596 | LYS539 | 97.75% | GLU592 | LYS539 | 96.25% |
| GLU543 | LYS728 | 88.50% | GLU543 | LYS728 | 96.00% |
| ASP569 | LYS415 | 73.00% | ARG232 | GLU176 | 84.75% |
| GLU592 | LYS539 | 44.75% | GLU596 | LYS539 | 77.75% |
| GLU307 | LYS253 | 43.50% | ASP569 | LYS415 | 65.50% |
| ARG588 | ASP620 | 17.00% | ARG235 | GLU176 | 15.00% |
| ARG541 | GLU592 | 15.00% | ARG588 | ASP620 | 5.00% |
| ARG541 | GLU596 | 0.50% | ARG130 | ASP519 | 3.00% |

| **Table S5. The details of interchain salt bridges of TYK2WT and TYK2V678F.** | | | | | |
| --- | --- | --- | --- | --- | --- |
| **TYK2 WT Dimer** | | | **TYK2 V678F Dimer** | | |
| **Residue1** | **Residue2** | **Occupancy** | **Residue1** | **Residue2** | **Occupancy** |
| ASP135 | LYS357 | 14.00% | GLU173 | LYS258 | 99.50% |
| ASP367 | ARG118 | 11.00% | ARG572 | ASP623 | 61.00% |
| ARG583 | GLU657 | 8.00% | ARG465 | GLU618 | 50.75% |
| ARG465 | GLU618 | 3.50% | ARG110 | GLU174 | 48.50% |
| GLU617 | ARG572 | 1.50% | ASP244 | LYS360 | 45.50% |
| ASP135 | ARG368 | 1.00% | ARG240 | GLU174 | 24.50% |
|  |  |  | ARG631 | ASP135 | 22.50% |
|  |  |  | ARG368 | GLU175 | 19.50% |
|  |  |  | ARG118 | GLU371 | 9.50% |
|  |  |  | ARG490 | GLU618 | 9.00% |
|  |  |  | ARG118 | GLU175 | 8.00% |
|  |  |  | ARG233 | GLU173 | 8.00% |
|  |  |  | ARG368 | GLU174 | 7.00% |
|  |  |  | ARG465 | GLU617 | 4.50% |
|  |  |  | ARG583 | GLU657 | 3.25% |
|  |  |  | ARG118 | GLU173 | 3.00% |
|  |  |  | ARG269 | GLU174 | 2.50% |
|  |  |  | ARG368 | GLU182 | 2.50% |
|  |  |  | ARG118 | GLU119 | 1.50% |
|  |  |  | ARG118 | GLU174 | 1.50% |
|  |  |  | ARG679 | ASP650 | 1.50% |
|  |  |  | ARG243 | GLU173 | 0.50% |
|  |  |  | ARG370 | GLU119 | 0.50% |
|  |  |  | ARG572 | ASP625 | 0.25% |

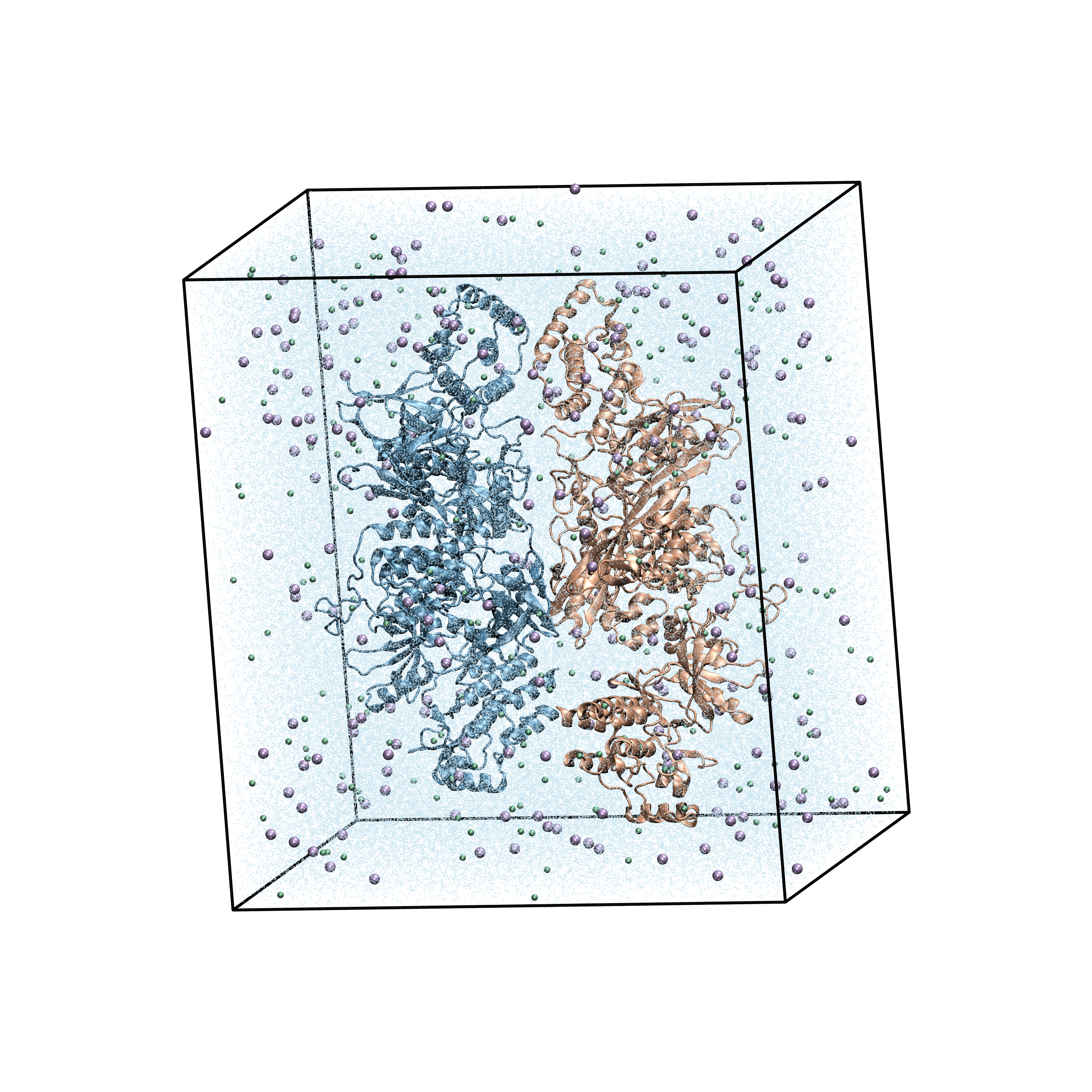

**Figure S1. Representative solvated simulation box.** The two protein monomers are shown as blue and orange cartoons, water molecules as light-blue dots, and ions as spheres. Black lines indicate the simulation box boundaries.

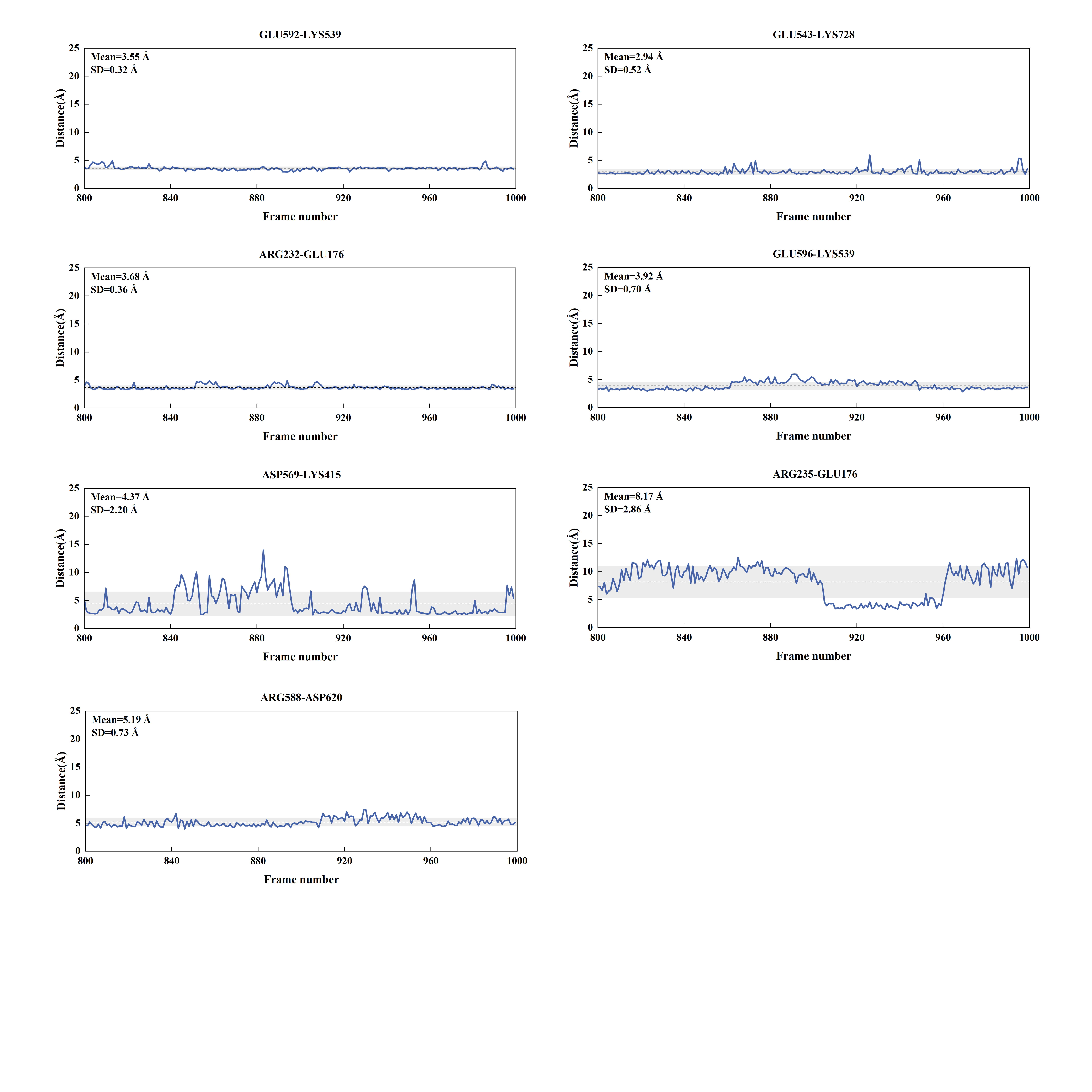
**Figure S2. Distances between atoms forming each salt bridge in the JAK2V617F dimer over the last 20 ns of simulation.** Each subplot shows the distance (y-axis) as a function of simulation frames (x-axis). The dashed line represents the mean distance, and the shaded area corresponds to the standard deviation.

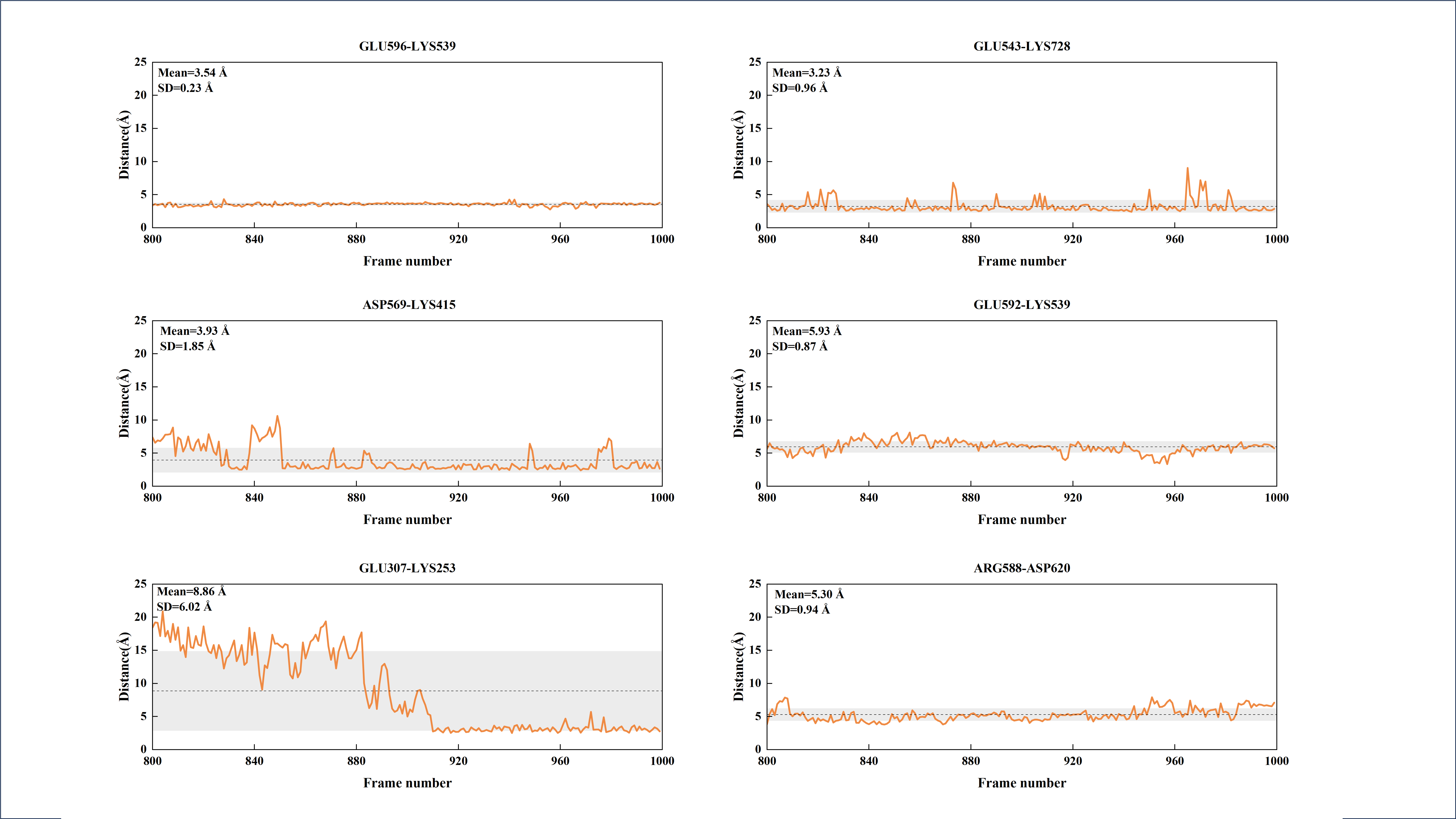

**Figure S3. Distances between atoms forming each salt bridge in the JAK2WT dimer over the last 20 ns of simulation.**

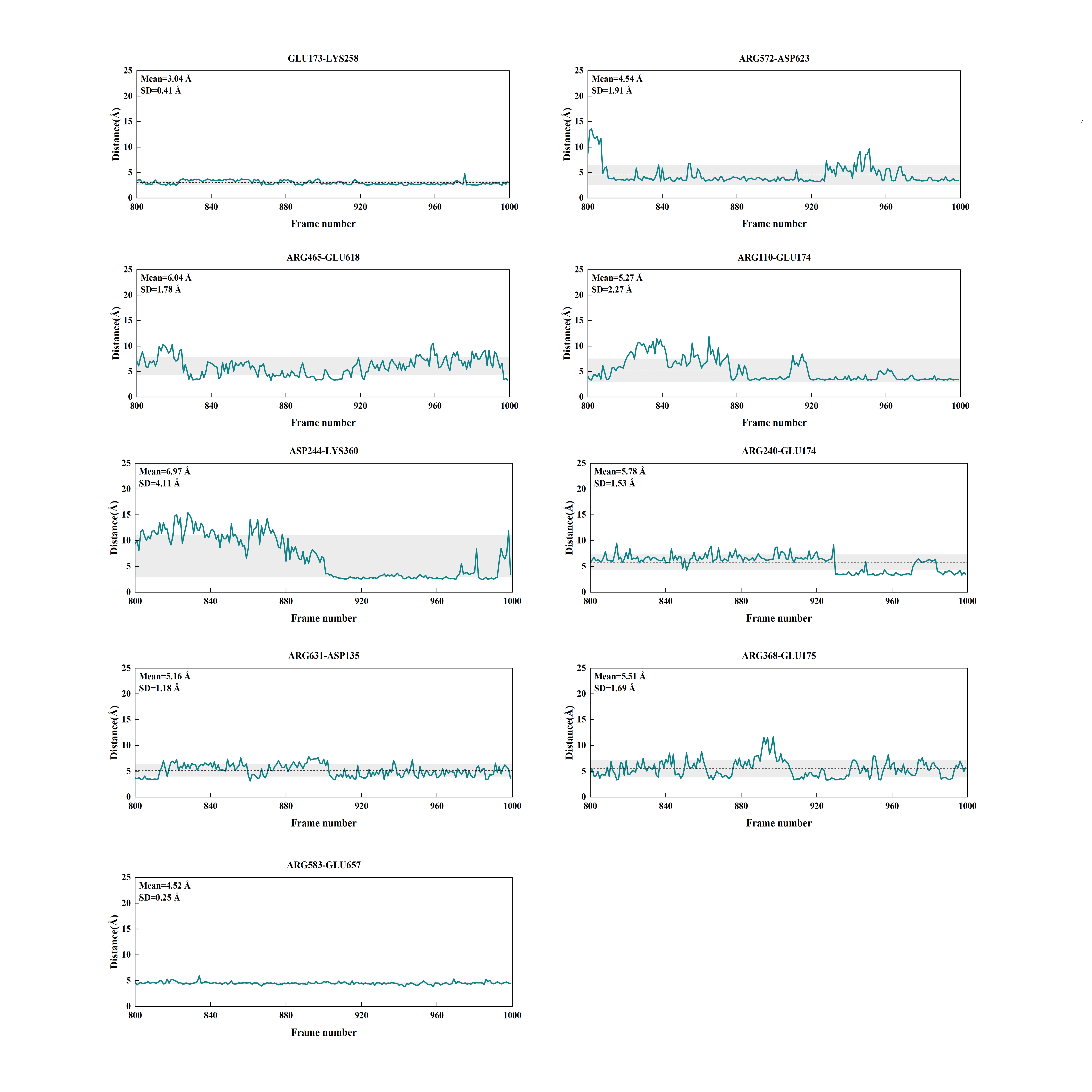

**Figure S4. Distances between atoms forming each salt bridge in the TYK2V678F dimer over the last 20 ns of simulation.**

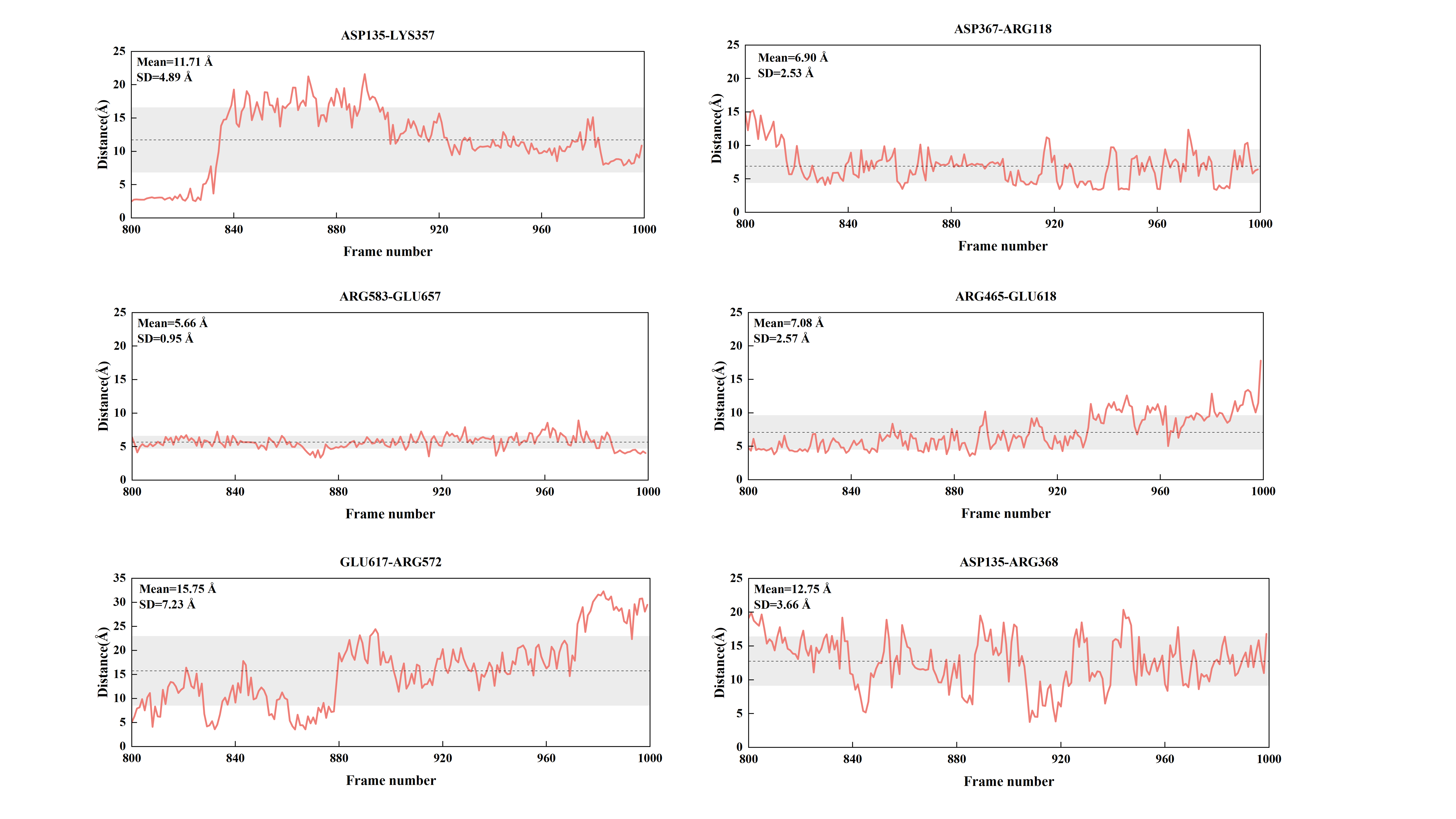

**Figure S5. Distances between atoms forming each salt bridge in the TYK2WT dimer over the last 20 ns of simulation.**
